# REPORTH: Determining orthologous locations of repetitive sequences between genomes

**DOI:** 10.1101/2024.10.14.618302

**Authors:** Prajwal Bharadwaj, Bram van Dijk, Frederic Bertels

## Abstract

Repetitive sequences are a common feature of bacterial genomes. Some repetitive sequences such as REPINs are mobile within the genome but inherited only vertically from mother to daughter across bacterial genomes. Selfish elements in contrast are mobile within the genome but also travel horizontally from genome to genome. Yet, no matter the nature of the association between the repetitive elements and the host, it is difficult to study the evolutionary dynamics of repetitive sequences across genomes. If it is unclear whether two sequences in two different genomes are in orthologous positions, it is difficult to infer parameters like the replication rate, horizontal transfer rate and rate of loss. Here we present a tool to facilitate these analyses called REPORTH. REPORTH determines whether repetitive sequences in different but closely related bacterial genomes occur in orthologous genomic positions. Whether a position is orthologous or not depends on the orthology of flanking sequences. Flanking sequences are deemed orthologous if they are bidirectional best hits. All repetitive sequences that are found in orthologous positions across different genomes are grouped together. Analyses of these groups can be used to study the evolutionary dynamics of selfish repetitive elements such as insertion sequences, but also for mutualistic repetitive elements such as REPINs.

## Introduction

Repetitive sequences are a common genomic feature. In eukaryotic genomes in particular, a large part of the genome is repetitive, largely because of the activity of transposable elements (Kidwell 2002). Bacterial genomes are much less repetitive, yet, there are many different repetitive sequence families in bacterial genomes (Haubold, Wiehe 2006; Silby *et al*. 2009). Broadly speaking there are mutualistic repetitive sequences such as REPINs (REP doublet forming hairpins) (Higgins *et al*. 1982; Bertels, Rainey 2011) or ribosomal DNA repeats (Lim *et al*. 2012) and selfish genetic elements such as insertion sequences, the most common transposable elements in bacteria (Lawrence *et al*. 1992; Mahillon, Chandler 1998).

REPINs are short repetitive sequences common in many bacterial genomes (Bertels, Rainey 2011). They are mobilized by a single copy transposase called RAYT (REP associated tyrosine transposase) (Nunvar *et al*. 2010; Ton-Hoang *et al*. 2012; Bertels, Gallie, *et al*. 2017). Unlike most other repetitive sequences in bacteria, REPINs are not horizontally but vertically inherited and hence are likely in a mutual relationship with their host (Park *et al*. 2021; Bertels, Rainey 2023). Because REPINs are vertically inherited all REPINs of the same type in a bacterial genome presumably descended from a common ancestor that either is still present in the genome (unlikely), or that was present in an ancestral strain but has been lost. The presence of REPIN parents, grandparents, siblings and cousins in the same genome allows the inference of parameters on the evolutionary dynamics of REPINs as, for example, REPIN duplication rate, which was estimated to be in the order of ∼10^−8^ duplications per REPIN and bacterial generation (Bertels, Gokhale, *et al*. 2017). However, it is still unclear how accurate these parameter estimates are, since the estimated parameters are based on the assumption that REPINs acquire mutations at neutral rates. Alternative measures that do not rely on sequence conservation could help us to understand duplication rates better.

In contrast to REPINs, insertion sequences travel horizontally rather than vertically through bacterial genomes (Bertels, Rainey 2023). Insertion sequences are also far better studied than REPINs across a wide variety of model systems (Sawyer *et al*. 1987; Lawrence *et al*. 1992; Mahillon, Chandler 1998). Since their duplication rates are magnitudes higher than that of REPINs (∼10^−5^ per element and bacterial generation), duplication rates can be measured in laboratory experiments (Sousa *et al*. 2013). Yet, studying the long term evolutionary dynamics of insertion sequences is similarly challenging. For example, knowledge about the presence of insertion sequences in orthologous extragenic spaces together with phylogenetic information can determine whether insertion sequences repeatedly insert into the same space or whether they are vertically inherited over long periods of time. Tracing the movement of insertion sequences within genomes as well as across genomes will certainly enhance our understanding of one of the most important bacterial parasites.

Distinguishing between orthologous and paralogous relationships is challenging even for non-repetitive gene families (Altenhoff, Dessimoz 2009). Orthologous groups of genes can be determined in various different ways. Simple approaches apply bidirectional best hits methods to determine orthologous genes in pairs of different species (Östlund *et al*. 2010), more complex approaches analyse best hit triangles (Tatusov *et al*. 1997), take gene trees into account (Wapinski *et al*. 2007) or analyse similarity networks of genes (Chen *et al*. 2006) to distinguish orthologous genes (similar as the result of speciation) from paralogous genes (similar as the result of gene duplication) (Fitch 1970; Koonin 2005). Distinguishing orthologous from paralogous genes is important, since orthologous genes often perform very similar functions while paralogous genes may have diversified to perform more different functions (Ohno 1970; Zeng *et al*. 2017). Functional similarities between orthologous genes has been used to automatically annotate genes belonging to the same orthologous group (Kuzniar *et al*. 2009). Functional gene annotations have later been used to understand and visualize synteny of genes across genomes with programs like WebFlaGs, Flankophile or full genome comparison tools like MAUVE (Darling *et al*. 2004; Saha *et al*. 2021; Thorn *et al*. 2023).

Functions of orthologous extragenic spaces can also be conserved (e.g. regulatory functions) and their evolution has been studied with programs like piggy (Thorpe *et al*. 2018). Here, we are mostly interested in extragenic spaces as containers for repetitive sequences (repeats). Comparing the locations of a sequence across different genomes can be used to distinguish between vertical and horizontal evolution hypotheses of the sequence. However, identifying the location of a sequence even between closely related strains can be difficult due to horizontal gene transfer and rearrangements (Raeside *et al*. 2014; Lee *et al*. 2016), which is why we have developed REPORTH.

REPORTH can identify the location of repeat sequences across closely related bacterial strains by determining orthology of flanking genes. REPORTH compares the flanking sequences of each repeat with the flanking sequences of all repeats in another genome. A bidirectional best hit strategy of the flanking sequences then allows the grouping of orthologous repeat sequences based on their location in the genome. As a proof of principle we have applied REPORTH to identify orthologous REPINs across 42 *Pseudomonas chlororaphis* strains, a bacterial species that contains three different REPIN-RAYT systems (Bertels, Rainey 2023). Identifying orthologous repeats can be used to measure duplication rates, loss rates, as well as repeat conservation and conversion studies across genomes.

## Methods

### REPORTH

To perform the clustering described here, a bioinformatics tool was developed, written in Python, called REPORTH (available at https://github.com/blackthorne18/reporth_cli). This tool can be installed using the PIP command *pip install reporth*. A detailed documentation is available at the GitHub repository. REPORTH requires BLAST+ (Camacho *et al*. 2009) to be installed and accesses BLAST functionality. All code and the dataset used in the analysis is available at https://github.com/blackthorne18/reporth_methods. The repository also contains README documentation required to replicate the results shown here. The clusters analysed here were generated using the command `reporth –-repin rarefan_output/ --genomes ./input/genomes –reptypes 0,1,2`

### Input data

REPORTH requires a set of closely related and fully sequenced bacterial genomes as input. It also requires the location of the repetitive sequences that are to be analysed. There are two possible ways of providing repetitive sequence locations. In case of REPIN sequences, we recommend using the location of a RAREFAN output folder (Fortmann-Grote *et al*. 2023) as input for REPORTH. It is also possible to simply provide a list of genome coordinates, for example when working with other types of repeat sequences (e.g. IS-elements), or REPINs identified with other pieces of software.

#### RAREFAN output as input data

When a RAREFAN output folder is provided as input, REPORTH will go through the output folder and find all genomes directories ([genome]_[x]/) where x is any digit signifying the type of REP. Using the ‘--`reptype`’ tag, the range of x can be specified as ‘--`reptype 1,2,3`’. Within these folders the locations listed in the files [genome]_[x].ss are parsed and only REPINs that are longer than 50bp are retained, excluding REP singlets – singlets can be included by providing a lower value using the ‘--`minrepinsize INTEGER`’ parameter. REPORTH will also ignore duplicate sequences.

#### List of genome locations as input

The format of the input files is explained in the README file of the REPORTH Github page but requires the name of the genome, the start and the end positions of each repetitive sequence, and the name of the repeat sequence type.

### REPORTH algorithm

#### Identifying flanking sequences of the repetitive sequences (Figure 1A)

For each repetitive region two 1000bp flanking sequences are selected. Each flanking sequence starts at a distance of 250bp from the repetitive element. The size of the flanking region was chosen so the flanking region encompasses at least one gene. As genes will be located at varying distances to the repetitive regions due to varying sizes of extragenic spaces we selected the flanking region at a distance of 250bp from the repetitive element to account for varying sizes of extragenic spaces. These parameters can be changed using the tags ‘`--win INTEGER`’ for window size, ‘--`fsize INTEGER`’ for the size of the flanking region.

**Figure 1.**
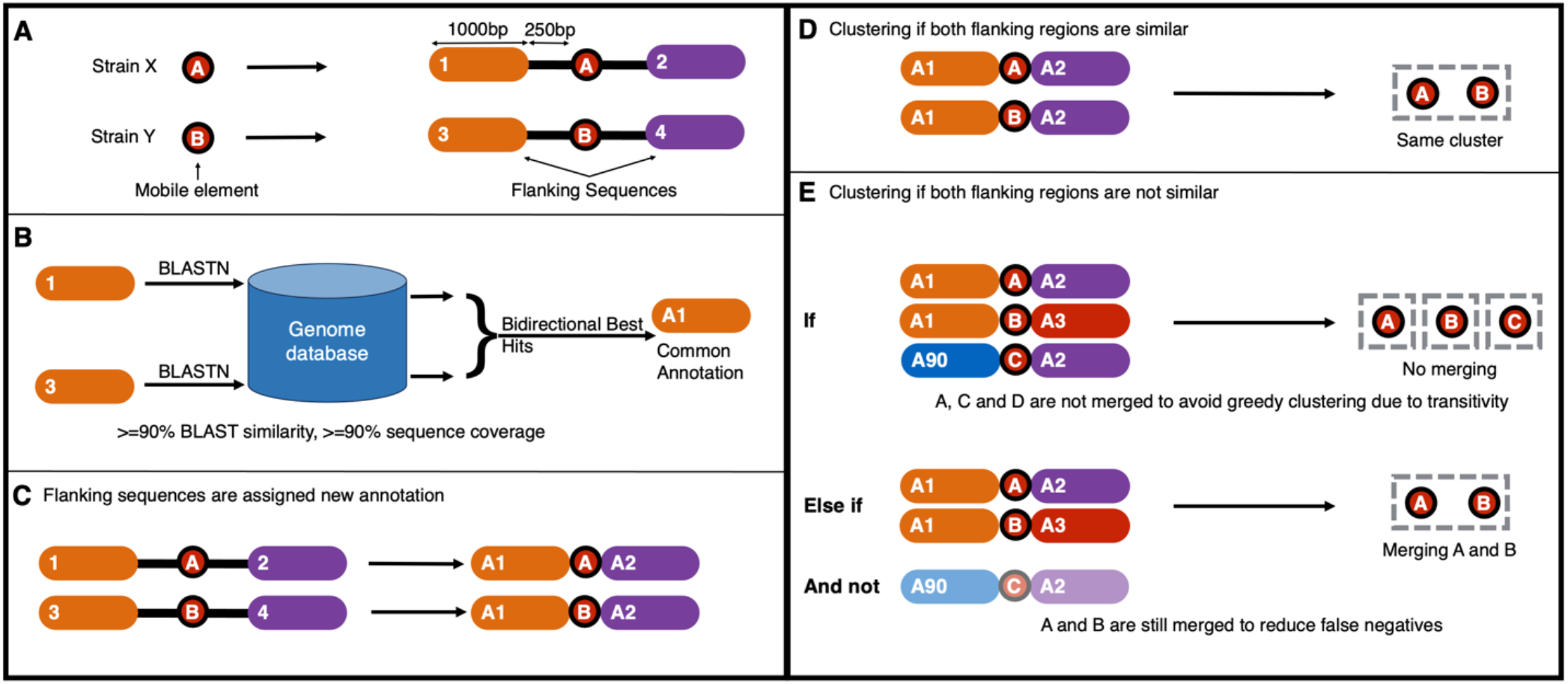
REPORTH workflow. (A) REPORTH needs a set of fully sequenced and closely related genomes as input. It also requires the location of repetitive sequences in these genomes. The locations can either be provided in the form of the location of RAREFAN output or a file listing the position of repetitive element in the respective genomes. Using this input and the provided genome files, a 1000bp flanking region on the 5’ and 3’ end of the repetitive element is selected at a distance of 250bp from the repetitive sequence. (B) For every repetitive element, BLASTN is used to identify homologs of both flanking sequences in all other genomes. (C) Two flanking sequences are bidirectional best hits of each other if each is the top ranking BLAST hit in the others genome. All bidirectional best hits are renamed to a common annotation. (D) Repetitive elements are remapped to have the new annotation of the flanking sequences associated with them. (E) All repetitive elements where both flanking sequences have the same annotation are grouped into a cluster (F) In the event that two repetitive elements only have one flanking sequence that matches, the decision of clustering them rests on the other flanking sequence of both repetitive elements. If either of the other (non-matching) flanking sequence have a common annotation with another flanking sequence, clustering is not performed.

#### Bidirectional best hits using BLASTN (Figure 1B and C)

The flanking sequences are searched against a database comprising all genomes provided by the user using BLASTN. For each flanking sequence queried, the top hit from each genome is selected only if the top hit itself overlaps at least 90% with the query and is identical across 90% of the query flanking sequence – which can be modified using the ‘--`pident INTEGER`’ and ‘--coverage INTEGER’ parameters. The query sequence and the top hit are classified as homologs if they form bidirectional best hits, meaning they are more similar in sequence to each other than any other sequence in their respective genomes. Bidirectional best hits have been shown to be reliable indicators of orthology (Östlund *et al*. 2010; Wolf, Koonin 2012).

#### Clustering of flanking sequences (Figure 1D)

Groups of sequences that formed bidirectional best hits are renamed to give a common annotation, yielding one or more orthologous groups. For example, if sequence 1 and sequence 2 formed bidirectional best hits, and the same sequence 1 formed a bidirectional best hit with sequence 3 from another genome, all three sequences would be renamed to, for example, “Sequence A”. The same is done until all sequences are assigned a label. We will define these groups of sequences as orthologous groups.

#### Assigning sequence clusters when both flanking regions are similar (Figure 1E)

All pairs of repetitive elements that are found in different genomes and flanked by orthologous sequences on both sides are assigned to the same repetitive sequence cluster.

#### Assigning sequence clusters when only one flanking sequence matches (Figure 1F)

If two repetitive sequences from different genomes only possess one pair of homologous flanking sequences, the decision of clustering them depends on the other flanking sequence of both repetitive elements. If either of the non-homologous flanking sequences are homologous to flanking sequences found in other sequence clusters, clustering is not performed. This prevents greedy clustering based on mobile elements moving in and out of flanking regions. We have also observed cases where neighbouring extragenic spaces were merged into a single extragenic space due to the deletion/disruption of the shared gene. Even in these cases we would consider the newly created space as separate from the ancestral extragenic spaces, and clustering is not performed. If the non-homologous flanking regions are not homologous to any other flanking regions, clustering is still performed.

### Visualisation and Output Files

REPORTH provides multiple output files to provide information on the clusters different repetitive sequences are sorted into and in order to understand why certain repetitive elements are clustered together.

**Table 1.** List of output files of REPORTH and a description of their contents.

| Output File | Description of contents |
| --- | --- |
| clusters_[date].txt | Listing all repetitive elements with unique cluster ID |
| meta_cluster_[date].txt | Lists the sequence similarity of the flanking sequences of a REPIN with the flanking sequences of every other REPIN in a cluster (in the order they are present in the cluster file) |
| lhs_hits.csv/rhs_hits.csv | CSV file storing the BLAST results of all flanking sequences |
| flanking_pairwise_dists.csv | Stores the pairwise distances between all flanking sequences from the BLAST hits |
| path_making_[date].txt | File listing whether a REPIN was merged into a cluster because of left flanking sequence/right flanking sequence or both |
| summary_histogram_1/2.png | Image file showing the distribution of number of cliques in a cluster. |

A graph showing the distribution of clique sizes of the clusters generated (summary_hisogram_1.png) is also provided which helps assess the precision of clustering of the entire dataset as a whole. Clusters having a single clique within them imply that the given cluster includes few outliers, while having two or more cliques likely implies that the region of the genome containing the repetitive element has evolved differently to its sister clade since the last common ancestor. More cliques imply higher divergence among the sequences. A summary_histogram_2.png file is also generated where it measures percentage similarity of flanking sequences in a cluster rather than clique sizes.

The path_making_[date].txt file shows the order in which REPINs were merged in a cluster. There are three headers in the file. Firstly, “>A_B with A_B both” indicates that all REPINs mentioned after this header were merged because both flanking sequences were homologous. Secondly, “>A_B with A_C” indicates that the REPINs that follow were merged because of only one flanking sequences. “A”, “B” and “C” are unique numerical identifiers that do not correlate with the final cluster number.

### Dataset

We tested REPORTH on a set of 42 genomes of *Pseudomonas chlororaphis*, which were obtained from The Pseudomonas Genome Database (Winsor *et al*. 2016). In these genomes REPIN sequences were identified using the RAREFAN (Fortmann-Grote *et al*. 2023) web service with the following settings: the query RAYT was yafM_SBW25, the ChPhzTR38 genome was used as the reference, the minimum seed occurrence was 55, distance group seeds was set to 15 and the e-value for identifying RAYT genes was set to 1e-30.

## Results

As a proof of concept we have applied REPORTH to REPIN-RAYT systems identified across 42 *Pseudomonas chlororaphis* genomes (Bertels, Rainey 2023). In bacteria there are a total of five different RAYT families (Bertels, Gallie, *et al*. 2017). Only two RAYT families are associated with REPINs: Group 2 and Group 3 RAYTs. *P. chlororaphis* contains three Group 3 REPIN-RAYT systems (and no Group 2 RAYTs), meaning that there are three different REPIN populations in each *P. chlororaphis* genome where each population is mobilized by a different Group 3 RAYT (Bertels, Rainey 2023). For ease of reference, we will refer to each of the populations by a color, either red, green, or blue.

We identified REPINs and RAYTs in each of the genomes using RAREFAN (Fortmann-Grote *et al*. 2022). REPIN sequences are two REP (**R**epetitive **E**xtragenic **P**alindromic) sequences in inverted orientation that appear within 200 base pairs of each other. RAREFAN identifies a total of 3005 blue, 5634 red and 6221 green REPINs across all genomes (**Figure 2A**). However, no genome contains more than 414 REPINs of any colour.

**Figure 2.**
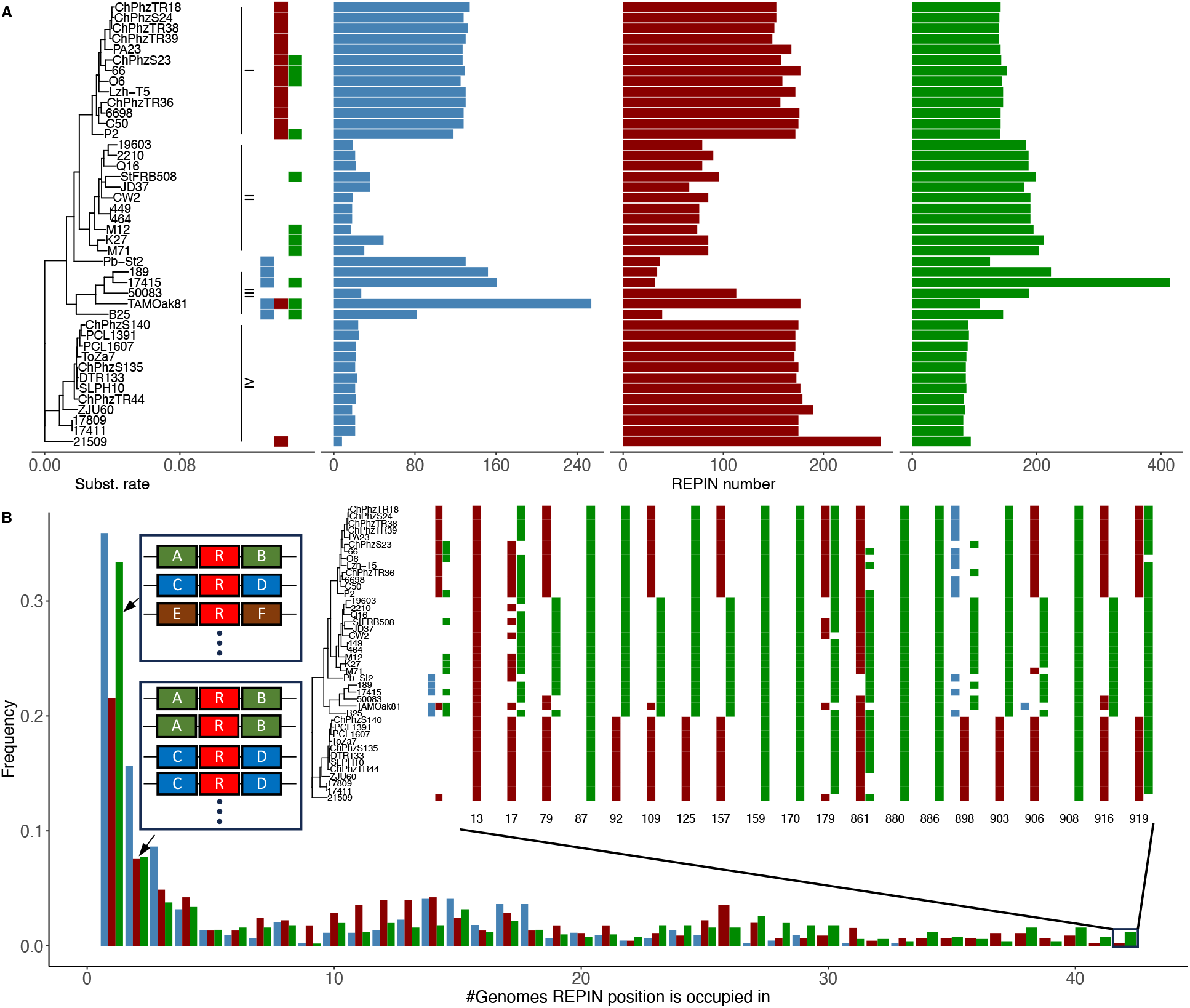
REPIN/RAYT distribution and cluster size distribution across *Pseudomonas chlororaphis*. (A) The left-hand side shows a phylogeny of 42 strains of *P. chlororaphis*, forming four distinct clades. Directly next to the phylogeny small bars indicate the presence of the red, blue, and green RAYTs in each of the *P. chlororaphis* strains. On the right side of the plot, the frequency of the three REPIN groups in each strain is shown. This figure was adapted from (Bertels, Rainey 2023). (B) The frequency distribution of cluster sizes across 42 *P. chlororaphis* strains is shown. The left side of the graph shows REPINs that are found in a specific extragenic space only once across all 42 strains. In the cartoon next to the bar chart “R”s indicate REPINs and [A-F] indicate different flanking sequences. The very right side of the bar chart shows REPINs that occur in an orthologous extragenic space across all 42 strains. The inset shows the REPIN composition of all 20 cases where an extragenic space is occupied by a REPIN in all 42 strains. Interestingly, only 7 out of 20 extragenic spaces are occupied by the same REPIN type. In 13 out of 20 cases a mix of REPINs is found in the space.

**Figure 2A** shows the REPIN/RAYT composition in 42 *P. chlororaphis* genomes that we have presented previously (Bertels, Rainey 2023). Red REPINs and RAYTs are predominantly found in clade I and clade IV. Green REPINs and RAYTs are predominantly found in clades I, II and III and blue REPINs and RAYTs in clade III. The REPIN distribution follows the distribution of the cognate RAYTs, except for blue REPINs that are abundant in clade I in the absence of blue RAYTs (Bertels, Rainey 2023). It is possible that either the red RAYTs in this clade evolved to mobilize blue REPINs or that the blue RAYT was lost in a recent common ancestor of clade I.

To get a fresh perspective on the data from **Figure 2A**, we can then directly apply REPORTH to the RAREFAN output. Across all 42 genomes, REPORTH identifies 1075 REPIN clusters. Each REPIN cluster contains REPINs that are flanked by orthologous sequences (i.e. based on bidirectional best hits, see **Methods**). The distribution of cluster sizes is shown in **Figure 2B**. Most clusters (253 out of 1075) only consist of a single REPIN sequence, which means there are no other REPINs in any of the other genomes that are flanked by a similar sequence. There are multiple possible reasons for this observation. Firstly, the extragenic space in which the REPIN is found may not exist in other genomes. Alternatively, the REPIN moved into this extragenic space recently and has not yet been duplicated. Thirdly, REPINs from this extragenic space were lost in related genomes. Finally, it is possible that the orthologous extragenic space in other genomes cannot accurately be identified.

On the other end of the spectrum, there are 20 REPIN clusters (1.9% of all clusters) that are consistently found across all 42 *P. chlororaphis* genomes (inset **Figure 2B**) in the same sequence context. In other words, these REPINs are found next to genes conserved across *P. chlororaphis*. There are six green REPINs that are completely conserved and one red REPIN (inset **Figure 2B**). There are no blue REPINs that are conserved across the entire species. Interestingly, 13 REPIN clusters are completely conserved but contain a mix of REPINs. Hence, the presence of a REPIN in the same extragenic space across different genomes does not mean that the REPIN sequence has been static, instead the data suggests that REPINs consistently present in conserved extragenic spaces change (they may be overwritten) as RAYTs are exchanged (gained, lost or horizontally transferred).

This preliminary analysis highlights the potential of REPORTH to provide insight into the biology and evolution of repetitive sequences. In the next section we will demonstrate that REPORTH results are reliable and consistent as long as input genomes are closely related.

### High similarity of flanking sequences in orthologous extragenic spaces

The average pairwise nucleotide distance between *P. chlororaphis* genomes does not exceed 4%. Vertically inherited genes should have diverged over a similar degree between genomes. Therefore, we defined that two flanking sequences are orthologous if they are at least 90%\ identical. This rule will therefore include genes even if they evolve more than twice as fast as the average gene in *P. chlororaphis*. Indeed, most flanking sequences (94%) share above 95% of their sequences, as expected from the analysis of genome wide pairwise similarities (**Figure 3**). Only about 0.4% of all flanking sequences are 90%-91% similar, indicating that a 90% threshold for sequence similarity should result in very few false negatives sequence clusters (<0.4%), sequence clusters that are missed because the flanking sequences have diverged too quickly during speciation. The proportion of false negatives is additionally reduced since clusters can also be formed if only one of the flanking sequences are more than 90% similar.

**Figure 3.**
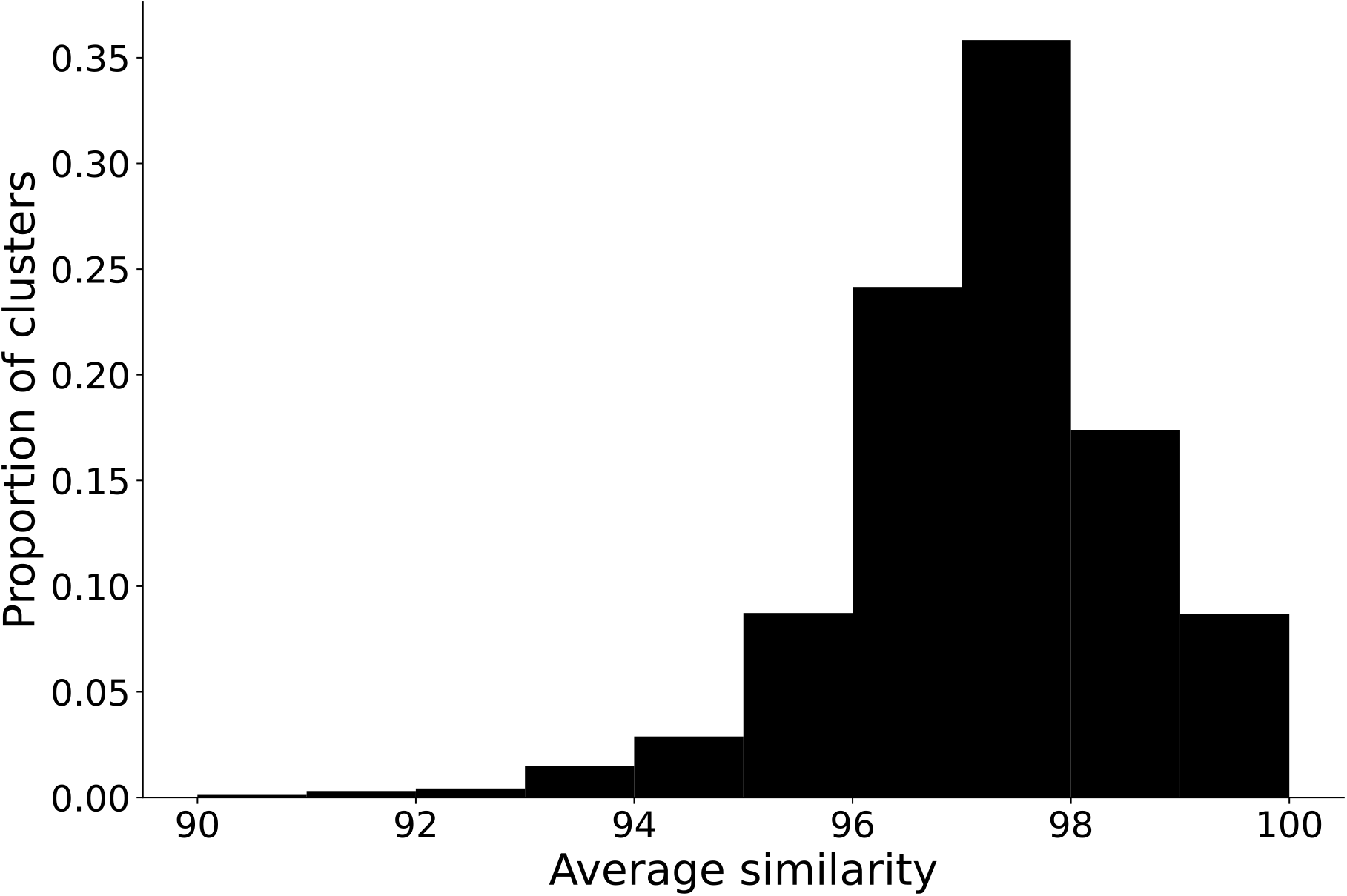
Average similarity in clusters. The figure shows the distribution of the average sequence similarity of both flanking sequences of REPINs present in the same cluster. For every cluster, the similarity of flanking sequences on one end and the flanking sequences on the other are taken separately, not averaged. The flanking sequences within a cluster are all highly similar to each other. In 96% of clusters, the sequence similarity of (both) flanking sequences of all REPINs within that cluster is greater than 86%. In only 4% of clusters, the sequence similarity is between 90-91%.

REPINs are clustered together when either both flanking sequences are orthologous, or when only a single flanking sequence is orthologous (>90% sequence similarity across more than 90% of the flanking sequence). As seen in **Figure 4**, the majority of REPINs (71%) are clustered together because *both* flanking sequences showed significant sequence similarities. About 29% of REPINs showed a one-sided sequence similarity. One-sided similarities are likely the result of a two-sided similarity being lost due to insertion of a mobile element, or due to a large chromosomal rearrangement.

**Figure 4.**
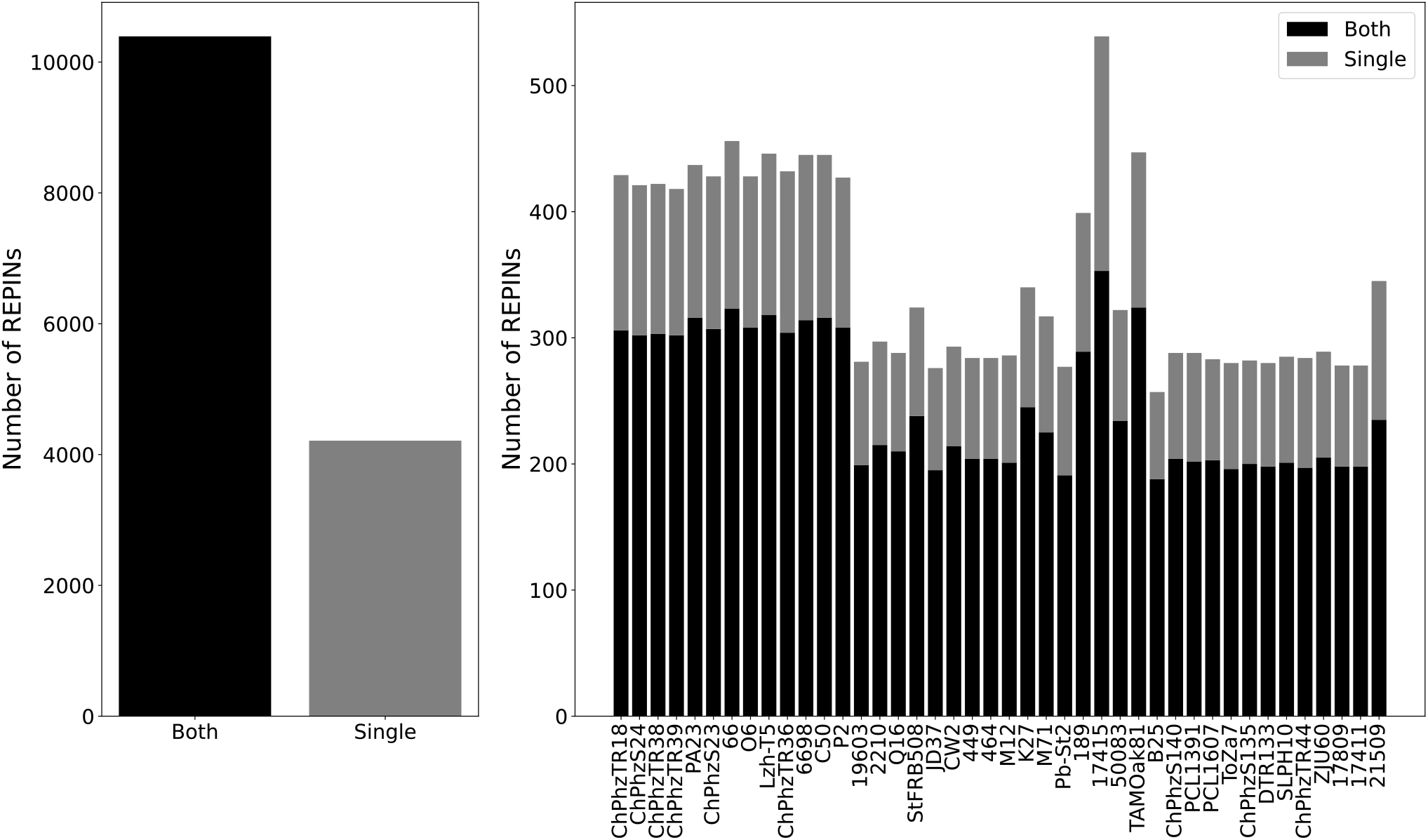
The basis for inclusion of REPINs in a cluster. (A) 71% of all REPINs were included in a cluster because both flanking sequences were orthologous with other REPINs from that cluster and 29% of them were included because only one of the flanking sequences was orthologous with others from the cluster (and the other non-matching flanking sequence had no other ortholog). (B) Number of REPINs from every genome that were included into a cluster based on similarity to both or a single flanking sequence(s).

### The presence of paralogous clusters is very rare

Paralogy, clustering sequences that are not derived from a recent common ancestor, cannot completely ruled out with bidirectional best hit methods (Wolf, Koonin 2012). Rare instances of recent gene duplication or horizontal gene transfer can lead to the clustering of paralogous instead of orthologous flanking sequences. It is possible to assess whether paralogy is a significant problem for our clustering method by determining all the number of flanking sequences where more than one sequence in a genome is more than 90% similar to the query sequence. Out of a total of 1075 sequence clusters we only identified 17 sequence clusters (1.6%) where incorrect clustering due to paralogy is possible (**Figure 5**). However, because sequences are clustered because of similarity of two flanking sequences the number of potential errors can be further reduced. Of all the cases where a flanking sequence has multiple hits within a genome, in only 4 out of the 17 clusters, was the flanking sequence that is a potential paralog used as a basis for merging the REPINs into a cluster. In the remaining 13 cases, the REPIN was merged into a cluster with both flanking sequences matching and one of those had a second copy in the genome. Hence the likelihood that REPORTH mistakenly clusters the wrong repetitive sequences due to multiple best hits (paralogy) is very low.

**Figure 5.**
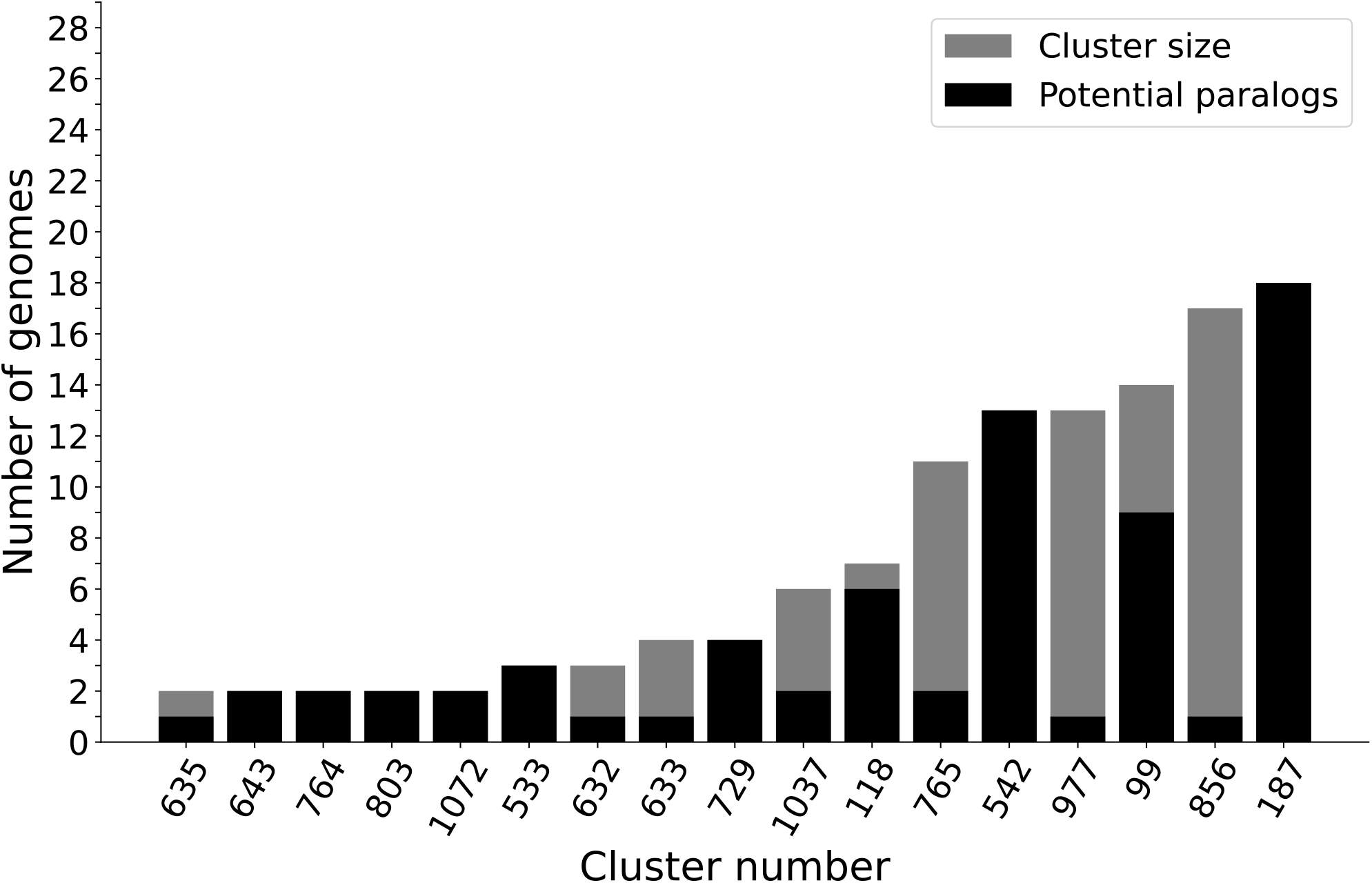
Presence of potential paralogs influencing clustering. The figure shows the proportion of REPINs within each cluster whose flanking sequence is a potential paralog (i.e. there is more than one homolog of that flanking sequence in that genome). The grey bar indicates the total number of REPINs in a cluster. The black bar shows the number of REPINs that were included into a cluster based on a flanking sequence that has multiple homologs in the genome.

## Discussion

Here we present REPORTH, a software tool that determines “orthologous positions” in the genome based on the orthology of flanking sequences. REPORTH is particularly useful for studying the evolution of repetitive sequences. It allows the comparison of repetitive sequences that are present in the same orthologous position across different genome. As a proof of principle we have applied REPORTH to a conserved class of repetitive sequences called REPINs (Bertels, Rainey 2011).

A distribution of cluster sizes first showed that REPINs are often found in unique positions in the genome, mostly because the orthologous extragenic space is not found in other genomes of the species. A significant proportion of REPINs is also conserved across the entire species. Interestingly, while REPINs are conserved in the same position, REPINs in that position change over time and correspond to the predominant RAYT gene (transposase responsible for REPIN movement (Nunvar *et al*. 2010; Bertels, Rainey 2011; Ton-Hoang *et al*. 2012)) in that genome. Our data suggests that RAYT genes can modify or replace REPINs inside the genome.

Further analyses of the REPIN data shows that REPORTH can robustly identify orthologous extragenic spaces. For closely related strains from a single species (>95% pairwise sequence similarity), our conservative threshold of 90% pairwise sequence identity allows the identification of robust sequence clusters. Requiring only one of the two flanking sequences to show high sequence similarity also allows us to identify sequences in conserved sequence spaces after genome rearrangements or the insertion of mobile genetic elements. This feature is particularly important because genome rearrangements are promoted by repetitive sequences due to homologous recombination (Consuegra *et al*. 2021) and as we show in our REPIN example, one sided orthology is surprisingly common. Finally, we show that distinguishing paralogy from orthology is not a big problem for REPORTH in our dataset of closely related bacterial genomes. Hence, as long as REPORTH is applied to closely related strains paralogy should not cause incorrect clustering.

Here we show how REPORTH can be used to study the evolution of REPINs. In the future we are planning to analyse REPIN clusters in detail to infer for example REPIN duplication rates as well as REPIN modification rates. However, REPORTH can also be applied to other types of repetitive sequences, such as insertion sequences (Mahillon, Chandler 1998). Preliminary analyses have shown that different types of insertion sequences can be distinguished by how frequently they are found in the same orthologous position in the genome. Suggesting that there are different insertion sequence “lifestyles” that lead to different sequence distributions.

Unfortunately, REPORTH can only be applied to very closely related sequences, which prevents the analyses of highly conserved repetitive sequences such as ribosomal RNA genes. If a similar program could be designed for distantly related genomes, then this could significantly enhance our understanding of rDNA or tRNA evolution. We could study how often these genes duplicate, how often they simply move to different positions in the genome and it would also facilitate studies of gene conversions.

Finally, we are convinced that REPORTH will facilitate the study of repetitive sequences in fully sequenced and closely related bacterial genomes. Preliminary analyses in REPINs and insertion sequences have already shown that there are significant knowledge gaps in the evolution of these sequence types that can be closed by applying REPORTH.

## Acknowledgements

PB, BvD and FB acknowledge generous core funding by the Max Planck Society.

